# Neutron crystallography reveals novel mechanisms used by *Pseudomonas aeruginosa* for host-cell binding

**DOI:** 10.1101/2021.09.24.461693

**Authors:** Lukas Gajdos, Matthew P. Blakeley, Michael Haertlein, V. Trevor Forsyth, Juliette M. Devos, Anne Imberty

**Affiliations:** Life Sciences Group, Institut Laue-Langevin, 71 Avenue des Martyrs, 38000 Grenoble, France; Partnership for Structural Biology (PSB), 71 Avenue des Martyrs, 38000 Grenoble, France; Univ. Grenoble Alpes, CNRS, CERMAV, 38000 Grenoble, France; Large Scale Structures Group, Institut Laue-Langevin, 71 Avenue des Martyrs, 38000 Grenoble, France; Faculty of Natural Sciences, Keele University, ST5 5BG Staffordshire, UK

**Author notes:** To whom correspondence should be addressed. Anne Imberty (, Twitter: @AnneImberty).

## Abstract

The opportunistic pathogen *Pseudomonas aeruginosa*, a major cause of nosocomial infections, uses carbohydrate-binding proteins (lectins) as part of its binding to host cells. The fucose-binding lectin, LecB, displays a unique carbohydrate-binding site that incorporates two closely located calcium ions bridging between the ligand and protein, providing specificity and unusually high affinity. Here, we investigate the mechanisms involved in binding based on neutron crystallography studies of a fully deuterated LecB/fucose/calcium complex. The neutron structure, which includes the positions of all the hydrogen atoms, reveals that the high affinity of binding may be related to the occurrence of a low barrier hydrogen bond induced by the proximity of the two calcium ions, the presence of coordination rings between the sugar, calcium and LecB, and the dynamic behaviour of bridging water molecules at room temperature. These key structural details may assist in the design of anti-adhesive compounds to combat multi-resistance bacterial infections.

## Introduction

The bacterium *Pseudomonas aeruginosa* is an opportunistic pathogen responsible for lung infections in cystic fibrosis patients and in immunocompromised patients, especially in hospital environments. It is particularly problematic in intensive care units. It is armed with an arsenal of virulence factors and antibiotic resistance determinants^1^ and has been identified as the number one priority, in the list of antibiotic-resistant bacteria by the WHO in 2017, in order to prioritize the efforts for the development of new antibiotics^2^. *P. aeruginosa* produces two soluble lectins^3,4^ LecA (PA-IL) and LecB (PA-IIL) specific for galactose and fucose, respectively, that are involved in the bacterial pathogenicity, adhesion and biofilm formation^5–7^. LecB is of special interest, since it is present only in a few opportunistic bacteria related to *Pseudomonas*^8^. The structure of LecB, and few related LecB-like, has a unique binding site with two closely located calcium ions (3.76 Å apart) that are directly involved in the sugar binding through the coordination of three hydroxyl groups of the carbohydrate ligand^9^ (Figure 1). The two calcium ions contribute to the receptor specificity since they coordinate monosaccharides with stereochemistry of two equatorial and one axial hydroxyl group present in fucose and mannose. These two calcium ions are also believed to play a role in the unusually strong affinity of LecB for fucose, which is in the low micromolar range, as compared to the millimolar affinity usually observed for lectin/monosaccharide interactions^10^. Carbohydrate chemists have proposed several fucose or mannose derivatives^11^ and some of these are able to inhibit biofilm growth, either as small molecules^12^ or as multivalent compounds^13^.

**Figure 1:**
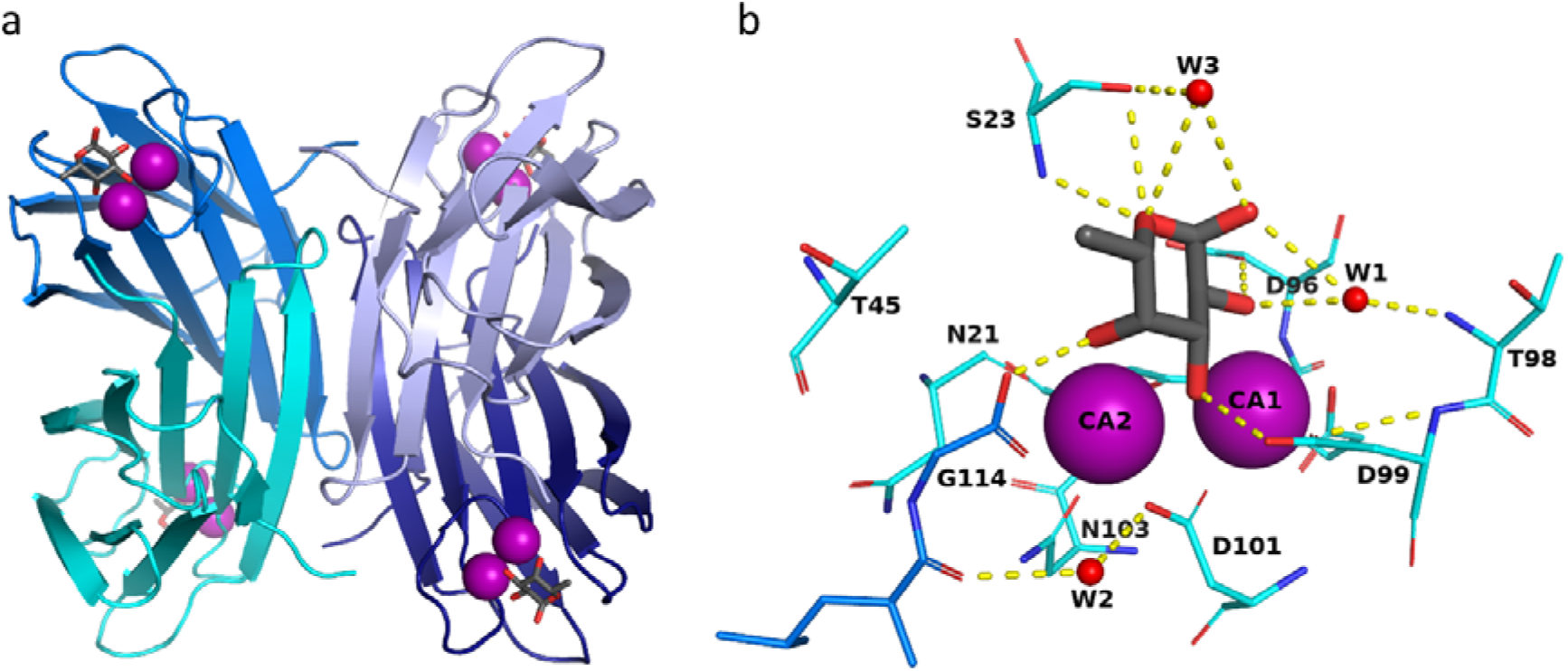
Structure of LecB lectin in complex with fucose. (a) Tetramer of LecB (PDB code: 1GZT). (b) Fucose binding site. Calcium ions and water molecules are represented by purple and red spheres respectively. Hydrogen bonds are shown as yellow dashed lines.

Neutron macromolecular crystallography (NMX) is the technique of choice for determining the position of hydrogen atoms and the protonation state of active groups^14–16^ that are involved in the catalytic activity of enzymes or the binding of a ligand to a macromolecule^17– 20^. Moreover, neutrons do not cause any radiation damage to the biological sample so the structure determination and data collection can be performed at room temperature (RT), giving a more accurate view of local flexibility and water mobility at biologically relevant temperatures^21,22^.

In the present work, perdeuterated fucose, produced through the use of engineered bacteria, has allowed, for the first time, the visualization of all hydrogen atoms (deuterium isotope) in the complex between a lectin from a human pathogen and its targeted sugar. We demonstrate the presence of several mechanisms underlying the observed high affinity binding, including a low barrier hydrogen bond, cooperative rings of hydrogen bonds and coordination contacts resulting in unique charge delocalization, and the mobility of trapped water molecules. Such structural details are of crucial importance for a more precise understanding of the protein/sugar/calcium triplet, and indeed, for the design of novel glycomimetic drugs that would reach higher affinity.

## Results and discussion

### Neutron structure of perdeuterated LecB/fucose complex

In order to determine the position of hydrogen atoms in the LecB/fucose complex, both molecules were first obtained in perdeuterated form. Hence, in the neutron analysis, hydrogen atoms are seen as deuterium, but for clarity in the following sections, are hereafter referred to as “hydrogen”. Recombinant perdeuterated LecB (D-LecB) was produced in an *Escherichia coli* high cell-density culture in the presence of D_2_O and glycerol-d_8_ after adaptation of cells^23^. Perdeuterated fucose (Fuc-d_12_) was obtained using recently described synthetic biology methods for which *E. coli* strains re-engineered for their carbohydrate metabolism were adapted for growth in deuterated media^24^. Co-crystals with volume of ∼0.1 mm^3^ were obtained over a period of approximately 4 weeks. The neutron structure of D-LecB complexed with Fuc-d_12_ was obtained at 1.90 Å resolution, and was jointly refined against both neutron and X-ray data (1.85 Å) collected from the same crystal at room temperature. The data collection and refinement statistics are shown in Table 1. As described previously, the overall fold of the LecB monomer is a β-sandwich composed of two curved sheets consisting of five and four antiparallel β-strands, respectively^25^ (Figure S1). Since the general features of the structure of LecB and the binding site with fucose have been well documented, the present work focuses solely on the position of hydrogen atoms and the effect of room temperature analysis.

**Table 1:**
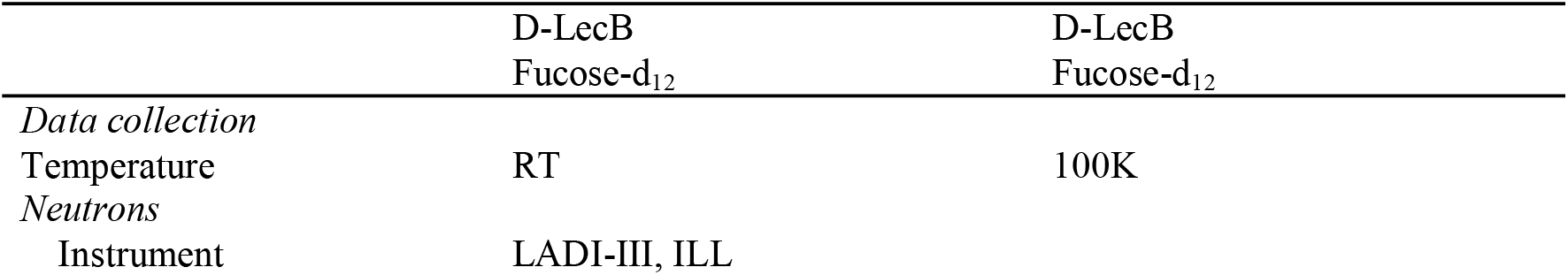
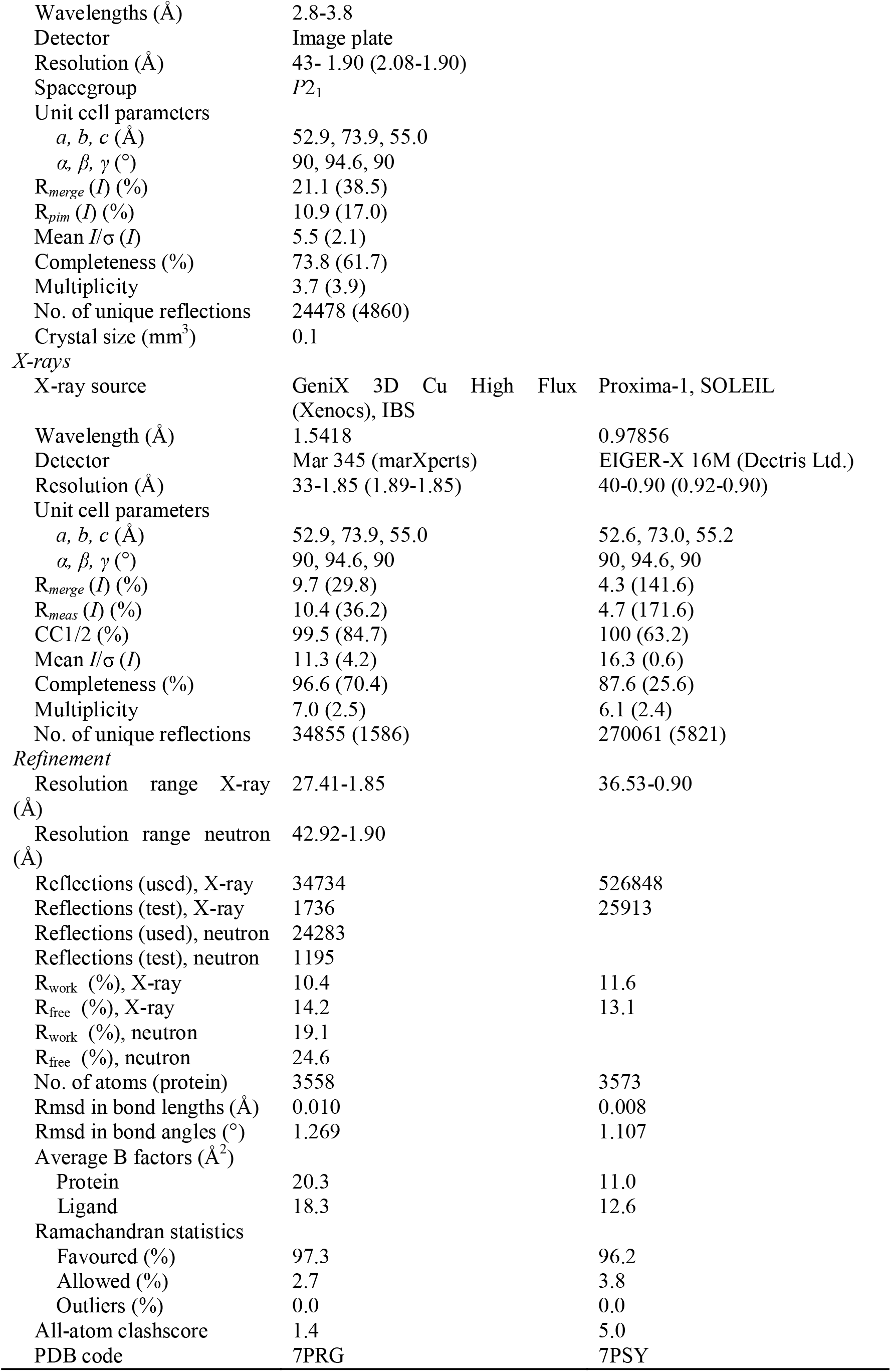
Room temperature neutron data collection and refinement statistics for the perdeuterated LecB/fucose complex

### Hydrogen atoms at monomers interface

The excellent quality of the neutron scattering length density maps (hereafter referred to as ‘neutron maps’) allows clear observation of the aliphatic amino acids and hydrogen bonds between adjacent β-sheets in the monomer. The tetramerization that occurs by the antiparallel association of β-strands from each dimer with their counterparts in the other dimer, involves mostly hydrogen bonds that could be directly visualised from the neutron maps. Furthermore, the use of perdeuterated protein has allowed the observation of the contacts involved in tetramer formation, such as the hydrogen bond between N-ter Ala1 of chain A and Asp75 of chain C, and the hydrophobic interactions between the methyl groups of the aliphatic amino acid side chains of Ala1 (chain A), Val77 (chain C) and Thr84 (chain D) (Figure S1b). The orientation of key water molecules and polar amino acid side chains could also be unambiguously determined from the neutron analysis (Figure S1c).

### Visualisation of hydrogen atoms on perdeuterated fucose

In all four chains, the fucose ring adopts the stable ^1^C_4_ chair conformation, with the α-configuration at the anomeric position. The use of perdeuterated fucose provided high-quality neutron density maps showing positive peaks for the deuterium atoms on the ring carbon atoms, the fucose methyl group, and the four hydroxyl groups (Figure 2). The neutron map clearly shows the hydrophobic contacts between the methyl group on the C6 position of fucose and the -CH_2_ and methyl groups of Ser23 and Thr45, respectively (Figure 2a). The deuterium atoms in the -CH_3_ group create a triangular-shaped neutron density and are in the most stable staggered conformation with respect to the aliphatic deuterium atom on C5.

**Figure 2:**
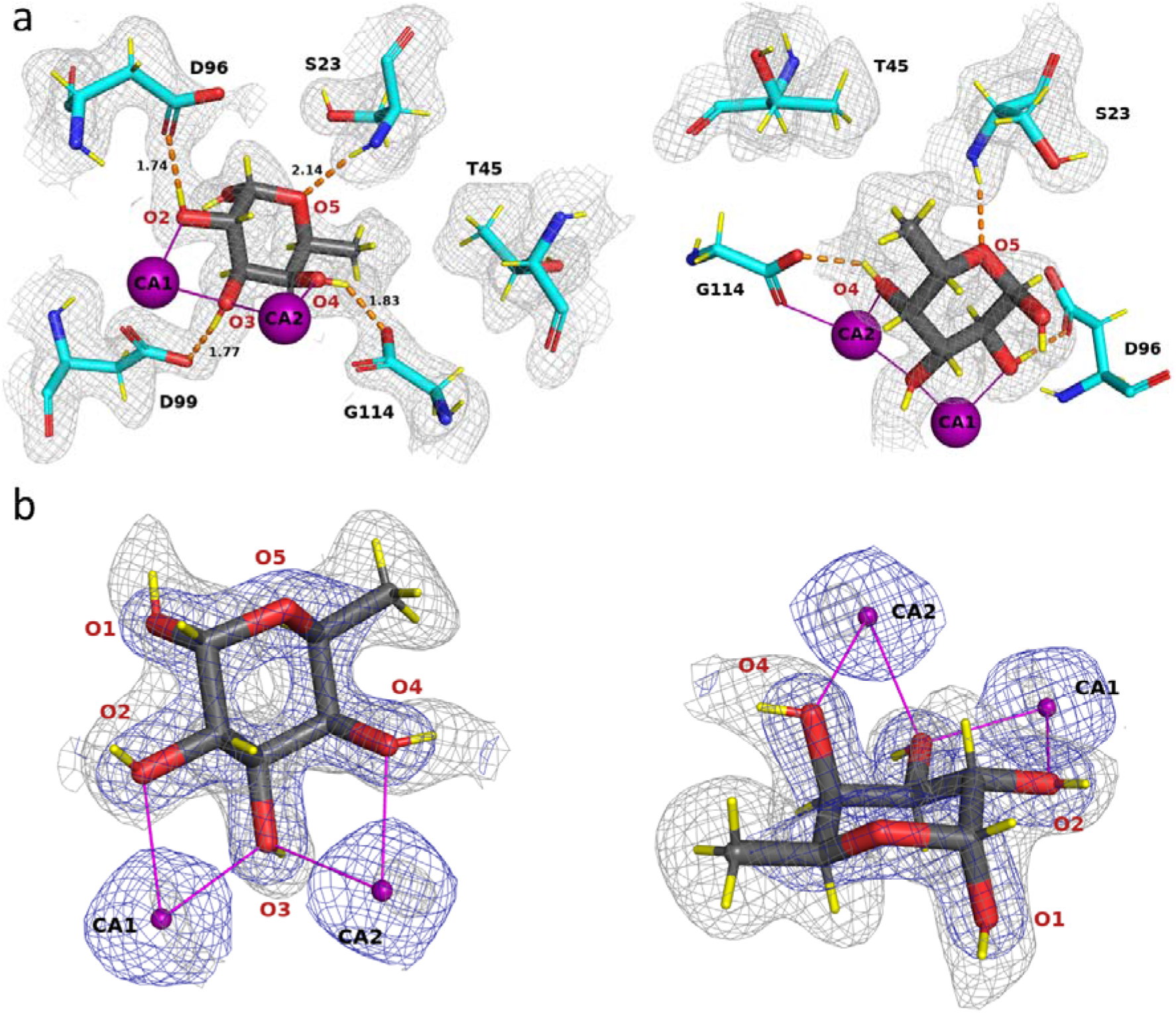
Fucose-binding site in the neutron structure of the D-LecB/Fuc-d_12_ complex. (a) Two different orientations of the same fucose-binding site (chain B). Fucose is represented as dark grey sticks and protein as cyan sticks. The calcium ions are shown as purple spheres and deuterium atoms are shown as yellow sticks. The hydrogen bonds and metal coordination are shown as orange dashed lines and as solid purple lines, respectively. The distances are given in Å. The 2m*F*_o_-D*F*_c_ neutron map (grey mesh) is contoured at 0.8σ. Amino acids involved in coordination of the calcium ions were omitted for clarity (Asn21, Asn103, Asp101, Asp104, Glu95). The first view (left) highlights the hydrogen-bonding network between fucose and the protein. The second view (right) shows the hydrophobic contact between methyl group of fucose and methyl and -CH_2_ groups of the Thr45 and Ser23 side chains, respectively. (b) The 2m*F*_o_-D*F*_c_ neutron map (grey mesh) and 2m*F*_o_-D*F*_c_ X-ray map (blue mesh), contoured at 1.4σ and 2.2σ, respectively.

The deuterium atoms attached to the four hydroxyl groups can be clearly visualized pointing away from the metal ions, establishing hydrogen bonds with the amino acid residues in the binding site. Peaks in the neutron density map can be observed for all of the hydrogen bonds established between the fucose and the protein (Figure 2 and Table S1). The Fuc-OD2 hydroxyl group donates its deuterium atom to OD1 of Asp96, Fuc-OD3 establishes a hydrogen bond with the side chain OD2 oxygen of Asp99, and Fuc-OD4 donates a deuterium atom to the OXT oxygen of the carboxyl group of Gly114 of the neighbouring monomer. The ring oxygen of fucose accepts a hydrogen bond from the amide backbone of Ser23.

For comparison, the 0.9 Å resolution X-ray structure, obtained at 100K, was also refined, showing the location of several hydroxyl groups in the m*F*_o_-DF_c_ omit map electron density (Figure S2). However, some hydrogens were not be located and the overall quantity of information is lower. Nevertheless, the analyses of the room temperature (neutron/X-ray) and 100K (X-ray) structures are also of interest for a comparison of thermal motion: In Figure S3 it is seen that two of the 4 monomers in the tetramer are more stable, and that there is less thermal motion than for the others. This observation may have implications for the hydrogen bonds within the active site and will be discussed later.

### Coordination of calcium ions

LecB and related lectins have unique carbohydrate receptors with two adjacent calcium ions in the binding site directly involved in sugar binding. Calcium atoms have a neutron scattering length of 4.70 fm, which is lower than that for C, O, N and D; hence their precise locations were refined from the X-ray data, confirming the close distance of 3.76 Å ± 0.02^10^. Each calcium is hepta-coordinated, with the involvement of three oxygen atoms of the fucose ligand (2 of these for each calcium ion), oxygen atoms from two asparagine residues and the carboxylate atoms of six acidic amino acids (three aspartates, one glutamate and the C-terminal group of Gly from the neighbouring monomer). The protonation states of these six acidic groups is a crucial question that can be best analysed from the neutron density map of the fully-deuterated complex that was used in this study. No hydrogen atoms could be detected on the acidic groups of the binding site, indicating that these are all negatively charged (Figure 3). The resulting total excess of 6 electrons is not fully compensated by the charge on the calcium ions and therefore the binding pocket of LecB has a strong negative charge.

**Figure 3:**
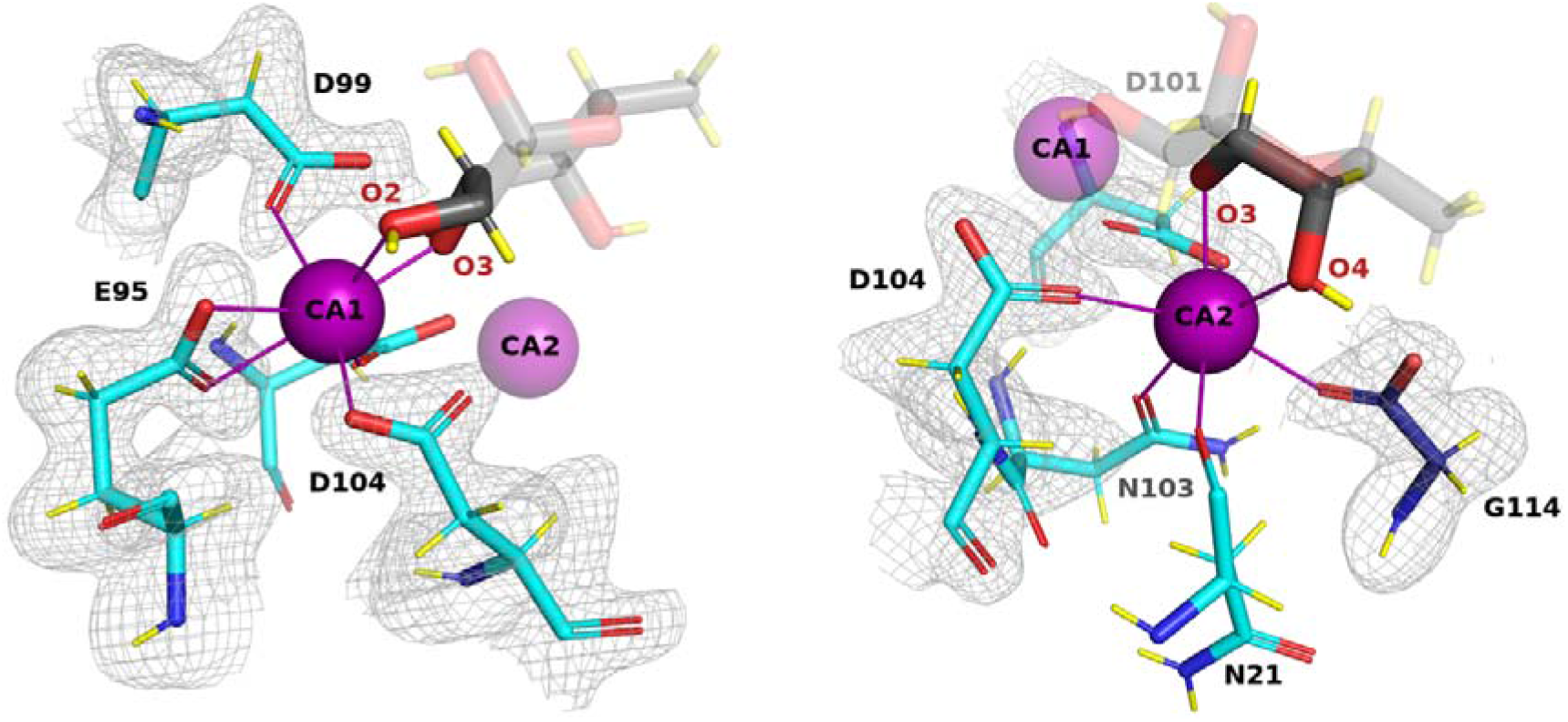
Calcium ions environment and protonation state of acidic residues. The 2m*F*_o_-D*F*_c_ neutron scattering length density (grey mesh) is contoured at 0.8σ. Calcium ions are shown as purple spheres and metal coordination as solid purple lines (binding site in chain B is shown).

### Effect of deuteration on structures and thermodynamics

The structures of the perdeuterated LecB/fucose complex can be directly compared to the previously reported hydrogenated LecB/Fuc complexes^26^ since the crystals were obtained using similar crystallization conditions. The structures are very similar with an RMSD of 0.13 Å over the whole tetramer main chain. The thermodynamics of fucose binding to LecB was also compared between the fully hydrogenated and fully deuterated systems, in H_2_O and D_2_O, respectively (Figure S4). No significant differences could be observed between the two studies with an enthalpy of binding ΔH of −30.45 (± 0.05) kJ/mol for the hydrogenated system and −30.95 (± 0.15) kJ/mol for the deuterated one. The dissociation constants were also almost identical with Kd values of 7.64 (± 0.15) μM and 7.96 (± 0.42) μM, respectively. Assays with mixing molecules (deuterated protein with hydrogenated ligand, or vice versa – data not shown) were also carried out and yielded similar results. This confirms that, for this particular protein/carbohydrate complex, the deuterated system is closely isomorphous with the unlabelled one, and that the results obtained for the visualization and the location of the deuterium atoms can be safely extrapolated to hydrogen atoms.

### Mechanisms for high affinity as inferred from neutron structure

The analysis of the neutron structure shown above has brought new insights to the mechanistic basis underlying the unusually high affinity of LecB towards fucose – providing information that could not have been obtained from X-ray diffraction data alone.

### Role of calcium ions in promoting a low-barrier hydrogen bond

Among the four hydrogen bonds between fucose and LecB, the three involving the carboxylate groups of amino acids coordinating fucose can be described as short (donor-acceptor distance < 2.6 Å) while the fourth one from the Ser23 main chain amine has a more standard length (distance > 2.9 Å)^27^. The nature of the short H-bonds was further investigated in order to determine the location of the deuterium atoms while removing the geometric constraints imposed during structure refinement. The m*F*_o_-D*F*_c_ neutron density maps, for which the deuterium atoms of the fucose hydroxyl groups were omitted, were therefore used to locate the peaks of deuterium atoms for chains A and D that display lower B-factors (Figure S3) and better quality maps (Table S2 and Figure 4A).

**Figure 4:**
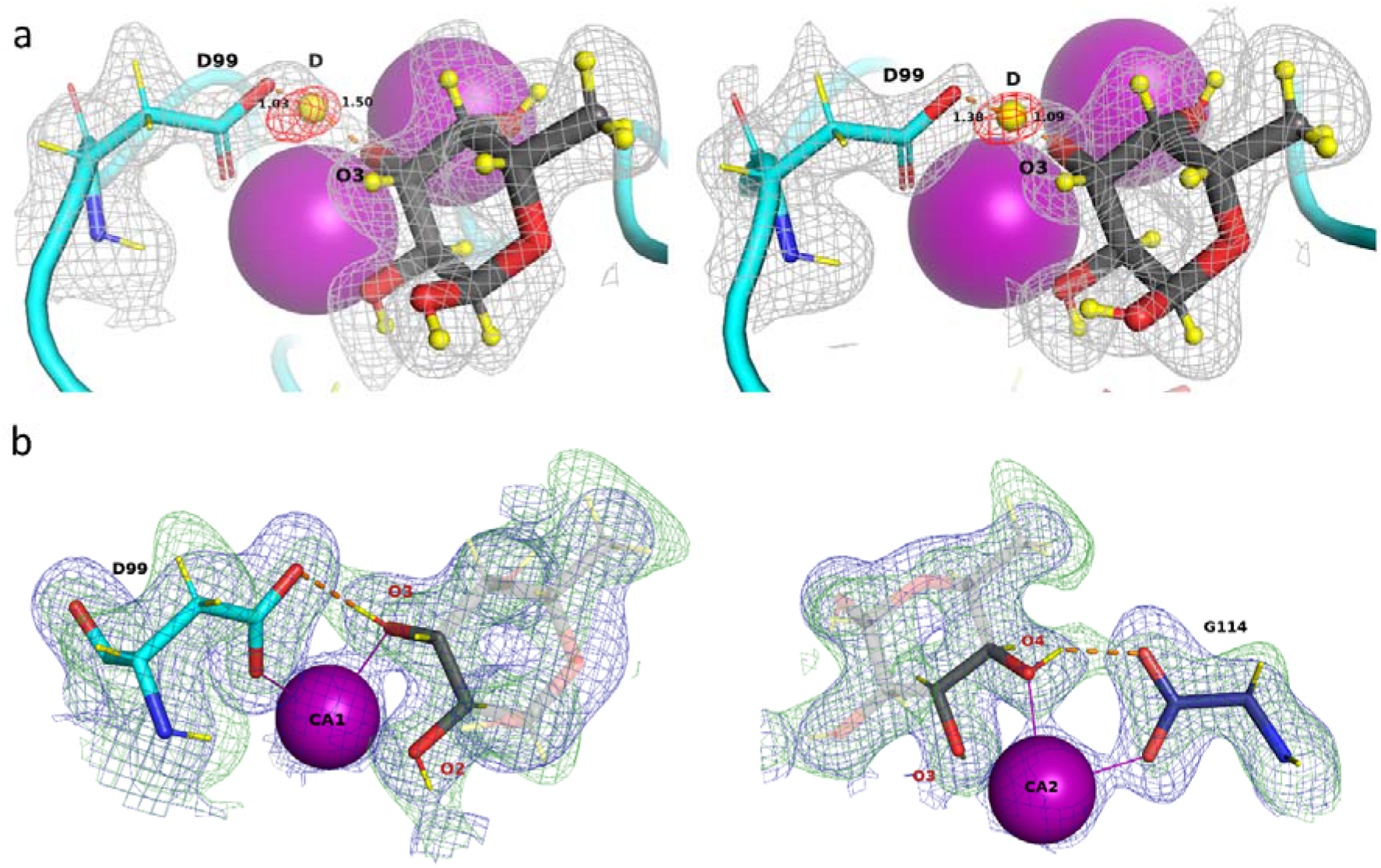
Mechanistic details of the synergistic interactions involving hydrogen bonds and metal coordination. (a) A low barrier hydrogen bond in chain A (left) and D (right) located from the omit difference m*F*_o_-D*F*_c_ neutron map (red mesh) in the vicinity of O3 of fucose. (b) Two six-membered rings formed by synergistic interactions between fucose, calcium and LecB. Hydrogen bonds are shown as orange dashed lines and metal coordination as solid purple lines. Deuterium atoms are shown as yellow spheres and yellow sticks. The 2m*F*_o_-D*F*_c_ neutron map (green mesh) is contoured at 1σ. The 2m*F*_o_-D*F*_c_ X-ray map (blue mesh) is contoured at 1.2σ.

The positive density peaks in which deuterium atoms can be placed are close to oxygens O2 and O4, and would correspond to short O-D distances of 0.85 ± 0.09 Å and 0.71 ± 0.01 Å, respectively, confirming a covalent link between deuterium and fucose oxygens. The resulting hydrogen bonds, involving Fuc-O2/Asp96 and FucO4/Gly114, can therefore be described as very short but classical. In contrast, the hydrogen bond between Fuc-O3D and Asp99 shows a peculiar character with a deuterium atom located almost equidistantly between the carboxylic side chain of Asp99 and the O3 atom of the fucose molecule (Figure 4a and Table S2) at 1.30 ± 0.21 Å away from O3 of fucose and 1.25 ± 0.17 Å OD2 of Asp99 with a very open O..D..O angle of 162.2 ± 1.8 °. The very short distance between the donor and acceptor (2.47 ± 0.04 Å) and, importantly, the delocalization of the hydrogen atom equally shared by two heteroatoms defines a low-barrier hydrogen bond^28–30^. Neutron maps are the most appropriate tool to visualise such bonds that have been observed not only in enzyme transition states, but also in protein-ligand complexes, such as in periplasmic phosphate binding protein^31^. The hydroxyl O3 of fucose is located at a very special location between the two calcium ions, with the deuterium pointing out towards Asp99; it is clear that the proximity of the two cations participates in its formation. It has been proposed that such bonds may be approximately six times stronger than classical ones^32^.

### Occurrence of synergistic rings of contacts

Three fucose oxygen atoms coordinate two calcium ions, with O2 coordinating the first calcium, O3 both calcium ions, and O4 coordinating the second calcium (Figure 3). The fucose hydroxyl groups O2 and O4 form hydrogen bonds with the acidic groups of Asp96 and the C-terminus of Gly114 of the neighbouring chain, respectively. Both interactions can be defined as very short hydrogen bonds (dist= 2.54 ± 0.04 / 2.49 ± 0.01 Å)^27^ although with no evidence of the delocalisation of the deuterium atom. They participate strongly in generating the high binding affinity. Furthermore, the coordination of calcium and hydrogen bonds creates a network of synergistic contacts between two aspartate residues, each coordinating one calcium ion and accepting a hydrogen bond from a fucose hydroxyl group, forming two 6-membered rings (Figure 4b). Previous quantum mechanical studies have suggested that, in presence of the fucose ligand, the charges of calcium delocalize very efficiently^26^.

### Mobility of water network in ligand binding site

Several conserved water molecules are commonly observed in LecB/Fuc complexes (PDB: 1GZT, 1OXC, 1UZV) and have established the existence of hydrogen-bonding networks between the protein, the ligand and the other ordered waters^10,25,33^. In the neutron structure, the water networks in each of the four monomers display small differences (Figure 5 and Figure S5). The most conserved water molecule, W1, accepts a hydrogen bond from the amide backbone of Thr98 (2.07 Å ± 0.05 Å) in all monomers, but adopts different orientations and therefore donates hydrogen atoms to either oxygen O1 and O2 of fucose, or to both of them, depending on the chains. This water molecule has been considered as very stable on the basis of previous high-resolution X-ray studies^10^; however, the present neutron structure at room temperature demonstrates higher mobility.

**Figure 5:**
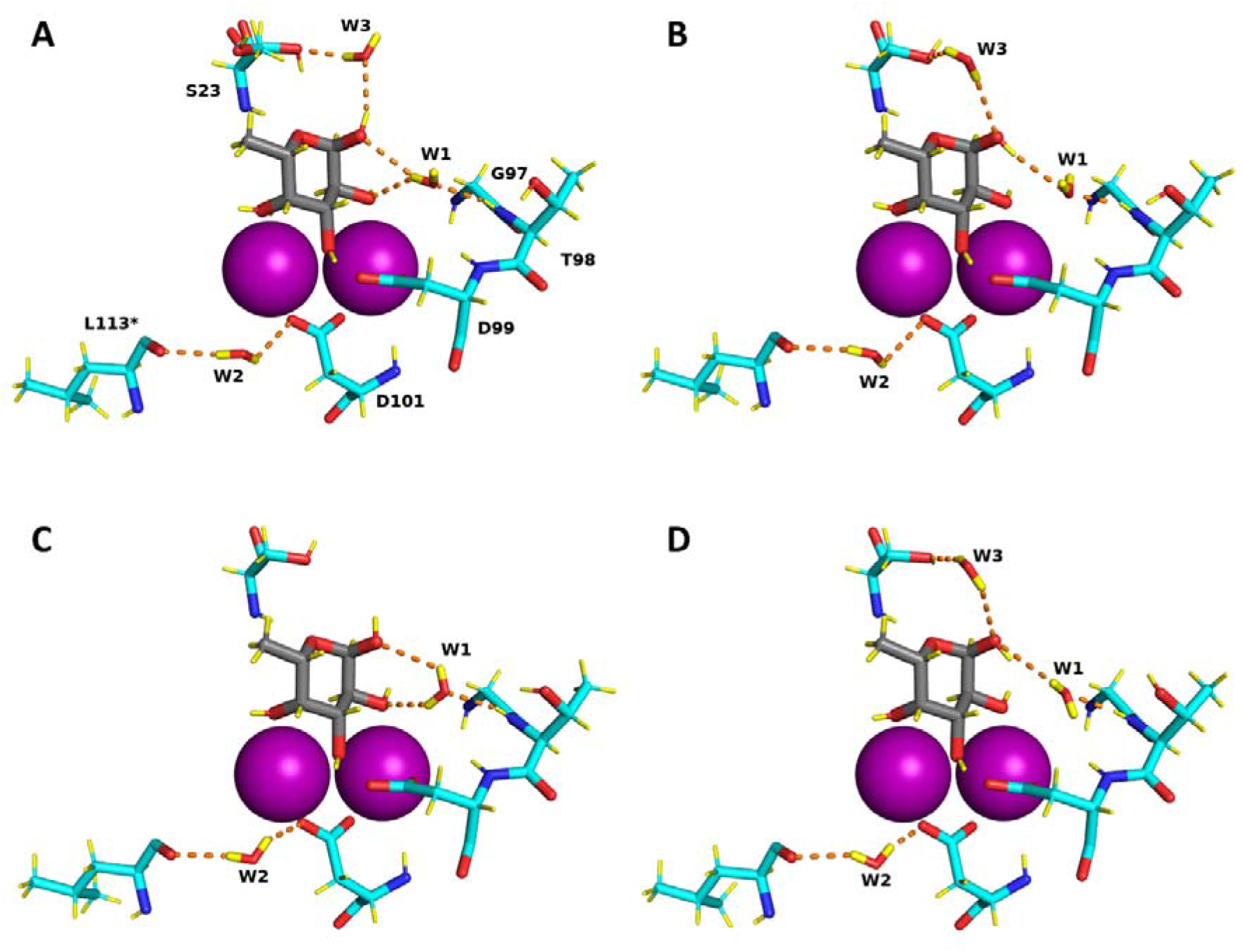
Water network in the fucose-binding sites in chains A, B, C and D of the perdeuterated LecB/Fuc-d_12_ complex determined from the neutron structure. Calcium ions are represented by purple spheres. Hydrogen bonds are shown as orange dashed lines. Water molecules in the binding sites are shown in stick representation.

This behaviour is not general. When examining another conserved water site, *i*.*e*. W2 bridging between Asp101 side chain and Leu113 main carbonyl group, this water is very stable and is implicated in similar hydrogen bonds for the four monomers. The third water molecule W3 bridges the anomeric oxygen atom to the side chain of Ser23 and is not conserved in all LecB crystal structures. In the neutron structure, W3 is present in three out of the four monomers. The neutron density is less clear for W3 but it could still be oriented based on the 2m*F*_o_-D*F*_c_ neutron map, as could the orientation of both OD1 fucose hydroxyl group and the side chain of Ser23.

The neutron structure therefore provides a different view from that provided by the previous high-resolution structures - allowing the discrimination of the fine details of the water networks in the four subunits of LecB, as well as showing disorder in the water molecules at the interface between carbohydrate ligand and protein. The binding of fucose therefore results in the release of water molecules strongly coordinated to the two calcium ions, and the protein/sugar interface does not create novel sites with strongly bound waters. While most protein-carbohydrate interactions display limited affinity because of unfavourable entropy barriers^34^, in this particular case, the overall entropy contribution is predicted to be favourable, in agreement with experimental measurements by ITC on this family of two-calcium containing lectins^35^.

## Conclusion

This is the first record of an experimental determination of the directionality of the fucose hydroxyl groups and the protonation state of acidic residues in the carbohydrate-binding site of LecB lectin from human pathogen *Pseudomonas aeruginosa*. Previously, a high-resolution 100K X-ray crystal structure (PDB: 1UZV) to 1 Å resolution allowed only one of the hydrogen atoms of O2-hydroxyl group to be oriented, pointing towards oxygen OD1 of the Asp96 side chain^10^. The protonation states of the acidic residues have not previously been determined and required us to use neutrons. The neutron analysis has provided evidence that all of the six acidic residues coordinating the calcium atoms are not protonated. This is in line with the prediction based on the *ab initio* calculations of bond distances in acidic residues having different protonation states^26^.

The unusually high affinity of LecB for fucose and fucosylated oligosaccharides can be rationalized by a synergy between short hydrogen bonds, including a low barrier one, and coordination bonds between the sugar, protein and calcium ions. Moreover, the binding of fucose causes displacement of three tightly-bound water molecules that are present in the sugar-free structure^36^ to the bulk solvent that is accompanied by a favourable entropy of binding. Both the high enthalpic contributions from strong hydrogen bonds, a favourable entropic term, together with charge delocalization caused by the involvement of two close calcium ions play a role in the high affinity. From room temperature neutron diffraction data it can be observed that water molecules which participate in the sugar binding display high mobility.

Medicinal chemistry, computer-based drug design or drug development, and in the future artificial intelligence structure based drug design all require accurate protein structures with accurate models, inclusive of hydrogen atoms. These will all benefit from high levels of structural detail of the binding sites in order to give the best possible results for drug development^37^. The information presented here will be important in rational structure-based drug design of novel potent inhibitors of *Pseudomonas aeruginosa*. It is believed that this work and these types of approaches will make important contributions to the development of glycomimetics that could help deal with the resurgence of multi-drug resistance bacteria.

## Materials and methods

### Protein expression

The LecB lectin was expressed in *Escherichia coli* BL21(DE3) bacteria harbouring a pET29b(+)-pa2l plasmid with a kanamycin-resistance gene. All cultures were grown at 37 °C with shaking at 180 rpm and were supplemented with 50 μg mL^-1^ kanamycin.

### Adaptation to D_2_O and deuterated glycerol-*d*_*8*_

Precultures of the LecB-producing strain were first grown in LB medium. Cells were then adapted to the deuterated Enfors minimal medium with the following composition: 6.86 g L^-1^ (NH_4_)_2_SO_4_, 1.56 g L^-1^ KH_2_PO_4_, 6.48 g L^-1^ Na_2_HPO_4_·2H_2_O, 0.49 g L^-1^ (NH_4_)_2_HC_6_H_5_O_7_ (diammonium hydrogen citrate), 0.25 g L^-1^ MgSO_4_·7H_2_O, with 1.0 mL L^-1^ of trace metal stock solution (0.5 g L^-1^ CaCl_2_·2H_2_O, 16.7 g L^-1^ FeCl_3_·6H_2_O, 0.18 g L^-1^ ZnSO_4_·7H_2_O, 0.16 g L^-1^ CuSO_4_·5H_2_O, 0.15 g L^-1^ MnSO_4_·4H_2_O, 0.18 g L^-1^ CoCl_2_·6H_2_O, 20.1 g L^-1^ EDTA), 5 g L^-1^ glycerol-d_8_ (Eurisotop). A single colony of *E. coli* containing the pET29b(+)-pa2l plasmid grown on an LB agar plate supplemented with 50 μg mL^-1^ kanamycin was used to inoculate 15 mL of LB medium with kanamycin. The preculture was then used to inoculate 15 mL of the hydrogenated Enfors minimal medium to OD_600_ of 0.1 and grown overnight. This preculture was used to inoculate 15 mL of the deuterated Enfors minimal medium to OD_600_ of 0.1 and grown overnight. This step was repeated 5 times prior to the fed-batch fermentation.

### Deuterated fed-batch fermentation of LecB

A final preculture of 150 mL was used to inoculate 1.2 L of deuterated Enfors minimal medium in a 3 L bioreactor (Infors, Switzerland). The pD of the culture medium was regulated at 7.2 by addition of 4% NaOD (Eurisotop, France). The temperature was maintained at 30 °C. After consumption of the deuterated glycerol-d_8_ from the culture medium, the fed-batch phase was initiated by continuous feeding with the additional 30 g of the deuterated glycerol-d_8_. The protein expression was induced overnight (17 h) with 1 mM isopropyl-β-D-thiogalactopyranoside (IPTG) at the OD_600_ of 13 at 30 °C. After the fermentation, the cells were recovered by centrifugation (10 500 g for 1 h at 6 °C) and the wet cell paste was frozen at −80 °C for long-term storage. The final yield was 51 g of the deuterated cell paste from a final volume of 1.6 L culture medium.

### Protein purification

Deuterated LecB was purified by affinity chromatography using the NGC system (Bio-Rad, Marnes-la-Coquette, France) on a 10 mL mannose-Sepharose resin packed in the C10/10 column. All buffers used for purification were prepared in H_2_O. Cells were resuspended in buffer A (20 mM Tris-HCl, pH 7.5, 100 mM NaCl, 100 μM CaCl_2_) in the presence of an EDTA-free protease inhibitor cocktail (cOmplete™, Roche) and treated with DENARASE® endonuclease (c-LEcta GMBH, Leipzig, Germany). Cells were lyzed using a cell disruptor with pressure of 1.8 kbar (Constant Systems Ltd., Northants UK). After centrifugation (24 000 x g for 30 min at 4 °C), the supernatant was filtered (0.45 μm) and the clear cell lysate was loaded onto 10 mL mannose-agarose column pre-equilibrated with the buffer A. After extensive washing the unbound proteins, the deuterated LecB was eluted with buffer A containing 100 mM free D-mannose. The purity of the protein was examined by 16% Tris-tricine SDS-PAGE stained with Coomassie Blue. The fractions containing the pure protein were pooled together and dialyzed against 5 L of buffer A at 4 °C for a week with changing the buffer once per day. The protein was concentrated using a 5 kDa cut-off concentrator (Corning® Spin-X® UF, England) and flash-frozen in liquid nitrogen for long term storage. The typical yield of the deuterated LecB was about 4 mg of protein per gram of wet cell paste.

### Isothermal titration calorimetry

All measurements were carried out at 25 °C using a MicroCal iTC200 isothermal titration calorimeter (Microcal-Malvern Panalytical, Orsay, France). Samples were dissolved in 20 mM Tris buffer, pH 7.5 with 100 mM NaCl and 100 μM CaCl_2_ prepared in H_2_O and D_2_O, respectively. A total of 20 injections of 2 μL of fucose (4 mM L-fuc/7.6 mM L-Fuc-d_12_) solution each were injected at intervals of 120 s with constant stirring of 750 rev min^-1^ in the 200 μL sample cell containing the LecB protein (0.46 mM LecB/ 0.50 mM D-LecB). Experimental data were fitted to the theoretical titration curve with the one set of sites model in the software supplied by MicroCal with ΔH (enthalpy change), Ka (association constant) and n (number of binding sites per monomer) as adjustable variables. The free energy change (ΔG) and entropy contributions (TΔS) were calculated from the equation ΔG= ΔH – TΔS = –RT ln *K*_a_ (T is the absolute temperature and R = 8.314 J K^-1^ mol^-1^). Two independent titrations were performed for each experiment.

### Crystallization

The perdeuterated LecB lectin was crystallized using a vapour-diffusion sitting drop method. The protein solution of 10 mg mL^-1^ was incubated with 250 μg mL^-1^ of perdeuterated fucose in the presence of 100 μM CaCl_2_ during 1 h at room temperature prior to crystallization. Perdeuterated fucose-d_12_ was produced by glyco-enginereed *E. coli* in a high cell-density culture as reported previously^24^. Co-crystals of deuterated LecB with L-fucose-d_12_ were obtained in the following conditions: 0.1 M Tris/DCl, pD 7.1, 20% (w/v) PEG 4000. The protein was mixed with the reservoir solution in 1:1 ratio and incubated at 19 °C.

### Neutron data collection and processing

Neutron quasi-Laue diffraction data from the crystal of perdeuterated LecB/fucose complex were collected at room temperature using the LADI-III diffractometer^38^ at the Institut Laue-Langevin (ILL) in Grenoble using a crystal of perdeuterated LecB in complex with perdeuterated fucose-d_12_ with volume of ∼ 0.1 mm^3^. A neutron wavelength range (Δλ/λ=30 %) of 2.8-3.8 Å was used for data collection with diffraction data extending to 1.9 Å resolution. The crystal was held stationary at different φ (vertical rotation axis) for each exposure. A total of 24 images were recorded (18 h per exposure) from 2 different crystal orientations. The neutron diffraction images were processed using the *LAUEGEN* software^39^. The *LSCALE* program^40^ was used to determine the wavelength normalization curve using intensities of symmetry-equivalent reflections measured at different wavelengths. The data were merged and scaled using *SCALA*^41^.

### X-ray data collection and processing

Room temperature X-ray data were collected from the same crystal used for the neutron data collection. The X-ray data were recorded on the GeniX 3D Cu High Flux diffractometer (Xenocs) at the Institut de Biologie Structurale (IBS) in Grenoble, France. The data were processed using the *iMosflm* software^42^, scaled and merged with *AIMLESS*^43^ and converted to structure factors using *TRUNCATE* in the *CCP4* suite^44^.

Other crystals of deuterated LecB co-crystallized with deuterated fucose were harvested manually and cryocooled at 100K by flash-cooling in liquid nitrogen. X-ray datasets were collected at the SOLEIL synchrotron (Saint Aubin, France) on the PROXIMA-1beamline. Images were recorded on the EIGER-X 16M detector (Dectris Ltd., Baden, Switzerland) and processed by *XDS*^45^.

### Joint neutron and X-ray refinement

The initial model (PDB: 1GZT) with water molecules, metal ions and ligands removed was used for the molecular replacement using *PHASER* in *PHENIX* suite. All further data analysis was done using the *PHENIX* package^46^. Refinement was performed using a restrained maximum-likelihood refinement in *phenix*.*refine*^47^. Restraint files for deuterated fucose were generated by *eLBOW* utility^48^ in *PHENIX*. Ligands were placed manually in *COOT*^49^. Water molecules were introduced automatically in *phenix*.*refine* based on the positive peaks in the m*F*_o_-D*F*_c_ neutron scattering length density and inspected manually. Deuterium atoms were introduced using the *Readyset* utility in *Phenix* and refined individually. All water molecules were modelled as D_2_O. Water oxygen atoms were initially placed according to the electron density peaks. The orientation of water molecules were refined and modified based on the potential hydrogen donor and acceptor orientation.

The quasi-Laue neutron R_work_ and R_free_ values for the final model were 19.1% and 24.6 %, respectively, while the X-ray R_work_ and R_free_ values were 10.4% and 14.2%, respectively. *Molprobity* software was used for structure validation. Refinement statistics are presented in Table 1. The models with the diffraction data have been deposited in the Protein Data Bank under accession codes 7PRG for the X-ray/neutron jointly refined model and 7PSY for the 0.9 Å 100K X-ray structure.

## Acknowledgements

The authors wish to acknowledge the ILL for provision of beamtime on LADI-III and technical support. We thank the ILL for the provision of studentship funding to L.G. V.T.F. acknowledges the UK Engineering and Physical Sciences Research Council EPSRC) for grants GR/R99393/01 and EP/C015452/1 that funded the creation of the Deuteration Laboratory within ILL’s Life Sciences Group. The authors acknowledge support from Glyco@Alps (ANR-15-IDEX02) and Labex Arcane/CBH-EUR-GS (ANR-17-EURE-0003). This work was supported by access to the HTX lab facility at EMBL and the PSB. We wish to acknowledge the IBS for access to the X-ray diffractometer and Proxima 1 beamline at SOLEIL Synchrotron, Saint Aubin, France for provision of beamtime. We wish to thank Annabelle Varrot and Sakonwan Kuhaudomlarp for their help in the X-ray data collection and Emilie Gillon for her help in the ITC measurements.

## Author contributions

A.I., M.H., J.M.D., M.P.B. and V.T.F. designed the experiment. L.G., J.M.D., M.H. and V.T.F. produced the deuterated biomolecules. L.G. purified the various components and carried out the experimental work. M.P.B. collected and processed the neutron data and provided expertise in structure determination. L.G. solved the crystal structures and refined them. L.G. prepared the figures and wrote the manuscript with A.I. and J.M.D., and with critical inputs from all authors.

## Competing interests

The authors declare no competing interests.

## Data availability

Coordinates and structure factors have been deposited in the Protein Data Bank under the accession codes: 7PRG, 7PSY.

## Notes

### Competing Interest Statement

The authors have declared no competing interest.

