## Supplemental information for "Neutron crystallography reveals novel mechanisms used by *Pseudomonas aeruginosa* for host-cell binding"

Supplementary Information

**Table S1: Geometrical parameters of the hydrogen-bonding network in the four binding sites of the LecB/Fuc-d_12_ complex in the neutron structure (with deuterium atom attached to donor with constrained distance of 0.85 Å during refinement procedure)**

| Donor | Acceptor | Hydrogen bond distance (Å)/ angle (°) | | | |
| --- | --- | --- | --- | --- | --- |
|  |  | Chain A | Chain B | Chain C | Chain D |
| *Direct interactions between the fucose residue and the protein* | | | | | |
| Fuc-O2 | Asp96.OD1 | 2.59 | 2.52 | 2.48 | 2.57 |
| Fuc-O3 | Asp99.OD2 | 2.51 | 2.53 | 2.49 | 2.43 |
| Fuc-O4 | Gly114.OXT^a^ | 2.49 | 2.47 | 2.48 | 2.51 |
| Ser23.N | Fuc-O5 | 2.92 | 2.96 | 2.94 | 2.96 |
| Fuc-OD2 | Asp96.OD1 | 1.81/165.1 | 1.74/179.1 | 1.77/151.5 | 1.86/162.6 |
| Fuc-OD3 | Asp99.OD2 | 1.74/160.0 | 1.77/161.8 | 1.92/133.3 | 1.69/153.7 |
| Fuc-OD4 | Gly114.OXT^a^ | 1.75/148.2 | 1.83/137.3 | 1.76/153.4 | 1.77/165.4 |
| Ser23.ND | Fuc-O5 | 2.10/169.8 | 2.14/169.1 | 2.12/176.4 | 2.16/171.2 |
| *Water-bridged hydrogen bonds between the fucose residue and the protein* | | | | | |
| Thr98.ND | O (Wat1) | 2.12/162.1 | 2.05/155.8 | 2.15/164.1 | 2.01/164.1 |
| Fuc-OD1 | O (Wat1) | - | 2.48/170.4 | - | 2.40/162.6 |
| D1 (Wat1) | Fuc-O1 | 2.46/144.4 | - | - | - |
| D1 (Wat1) | Fuc-O2 | 2.43/142.2 | - | 2.54/143.0 | - |
| D2 (Wat1) | Fuc-O1 | - | - | 2.56/125.6 | - |
| Fuc-OD1 | O (Wat2) | 2.24/160.8 | - | - | - |
| D2 (Wat2) | Ser23.OG | 1.87/149.5 | - | - | - |
| D1 (Wat2) | Fuc-O1 | - | - | - | 2.50/147.9 |
| D2 (Wat2) | Ser23.OG | - | - | - | 2.40/160.7 |

^a^ C-terminal residue from the neighbouring monomer

**Table S2: Geometrical parameters of the hydrogen-bonding network in binding sites of chains A and D with deuterium located in the omit map, with no constraints for O-D distance on fucose hydroxyl groups during refinement procedure. As defined in the results section on the importance of calcium, the hydrogen bonds involving** Fuc-O5/Ser23 is a normal hydrogen bond; Fuc-O2/Asp96 and FucO4/Gly114 are short but classical; Fuc-O3/Asp99 is considered a low-barrier hydrogen bond because of the position of the hydrogen equally shared by the two heteroatoms.

|  |  |  | Chain A | Chain D | average | St. Dev |
| --- | --- | --- | --- | --- | --- | --- |
| Dist donnor...acceptor (Å) | Fuc-O2 | Asp96.OD1 | 2.59 | 2.57 | 2.58 | 0.01 |
| O-D (Å) | Fuc-O2 | D | 0.94 | 0.76 | 0.85 | 0.09 |
| D….O (Å) | D | Asp96.OD1 | 1.68 | 1.86 | 1.77 | 0.09 |
| Angle O-D…O (°) | O2-D...Asp |  | 161.9 | 153.2 | 157.6 | 4.4 |
| Dist donnor...acceptor (Å) | Fuc-O3 | Asp99.OD2 | 2.51 | 2.43 | 2.47 | 0.04 |
| O-D (Å) | Fuc-O3 | D | 1.5 | 1.09 | 1.30 | 0.21 |
| D….O (Å) | D | Asp99.OD2 | 1.03 | 1.38 | 1.21 | 0.17 |
| Angle O-D…O (°) | O3-D...Asp |  | 164 | 160.4 | 162.2 | 1.80 |
| Dist donnor...acceptor (Å) | Fuc-O4 | Gly114.OXT | 2.49 | 2.51 | 2.50 | 0.01 |
| O-D (Å) | Fuc-O4 | D | 0.71 | 0.71 | 0.71 | 0.00 |
| D….O (Å) | D | Gly114.OXT | 2.2 | 1.91 | 2.06 | 0.15 |
| Angle O-D…O (°) | O4-D...Gly |  | 106.3 | 144.2 | 125.3 | 18.9 |
| Dist donnor...acceptor (Å) | Ser23.N | Fuc-O5 | 2.92 | 2.96 | 2.94 | 0.02 |
| O-D (Å) | Ser23.N | D | 1.06 | 0.81 | 0.94 | 0.13 |
| D….O (Å) | D | Fuc-O5 | 2.35 | 2.52 | 2.44 | 0.09 |
| Angle O-D…O (°) | Ser-D...O5 |  | 110.5 | 115.4 | 113.0 | 2.45 |

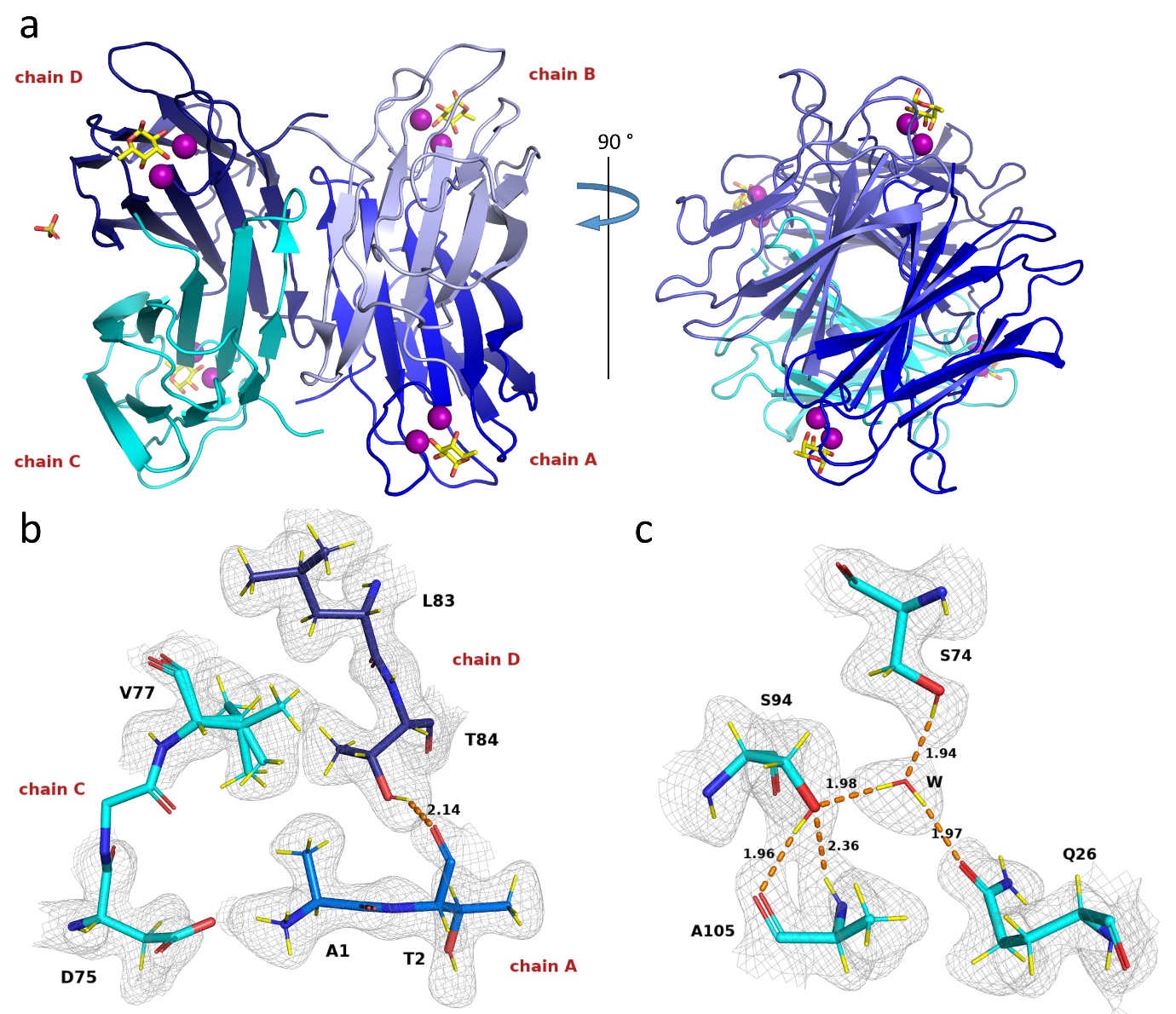

**Figure S1: Room temperature structure of D-LecB in complex with Fuc-d_12_.** Calcium ions are shown as purple spheres, fucose molecules are shown as yellow sticks, a sulphate ion is shown as orange and red sticks. Hydrogen bonds (Å) are shown as orange dashed lines. Deuterium atoms are coloured yellow (panels b and c). The 2m*F*_o_-D*F*_c_ neutron density (grey mesh) is contoured at 0.8σ. (a) Overall structure of D-LecB tetramer in complex with Fuc-d_12_. (b) Protein interface between chains A, C and D. (c) Hydrogen bonding network involving a water molecule.

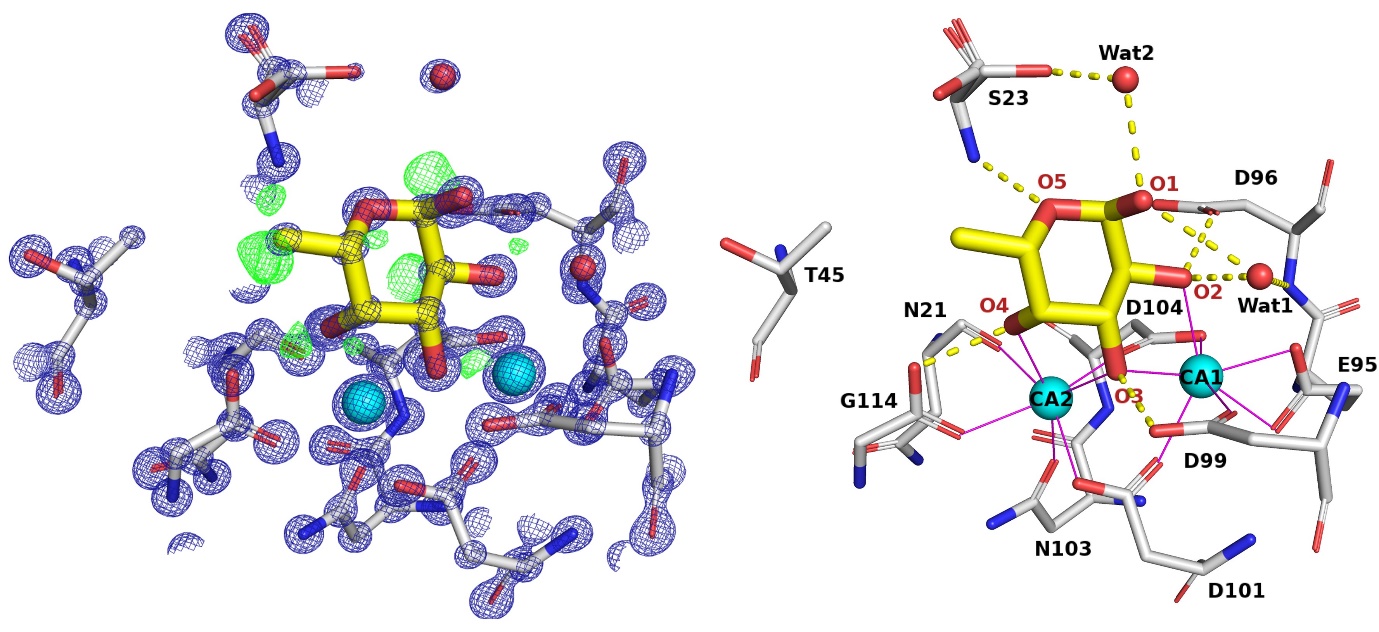

**Figure S2:** **Stick representation of the** **0.9 Å 100K X-ray structure of the fucose-binding site of the perdeuterated LecB/Fuc-d_12_ complex**. Fucose is shown as thick yellow sticks and the protein as thin grey sticks. 2m*F*_o_-D*F*_c_ electron density (blue mesh) is contoured at 4σ. The m*F*_o_-D*F*_c_ omit electron density (green mesh) is contoured at 2.5σ showing the positions of deuterium atoms on the fucose. Hydrogen bonds are shown as yellow dashed lines and distances are in Å. The metal coordination is represented by purple solid lines. The calcium ions are shown as cyan spheres.

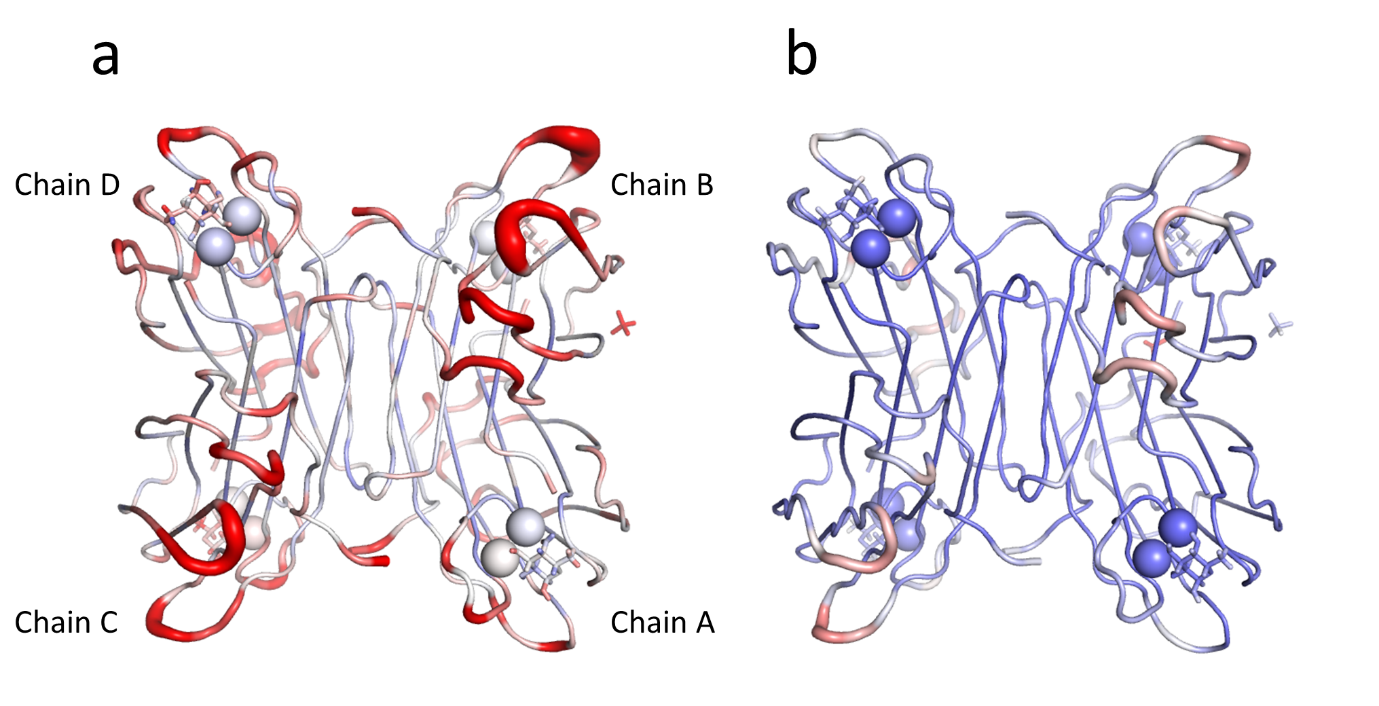

**Figure S3: Comparison of B-factors at room temperature (neutron/X-ray, left) and 100K structures (X-ray, right) of perdeuterated LecB/Fuc-d_12_ complex.** Graphical representation was performed with the “cartoon putty” option in Pymol with colour coding from 3 (blue) to 20 (red) for B-factor. Chains A and D display lower B-factors and therefore better quality maps.

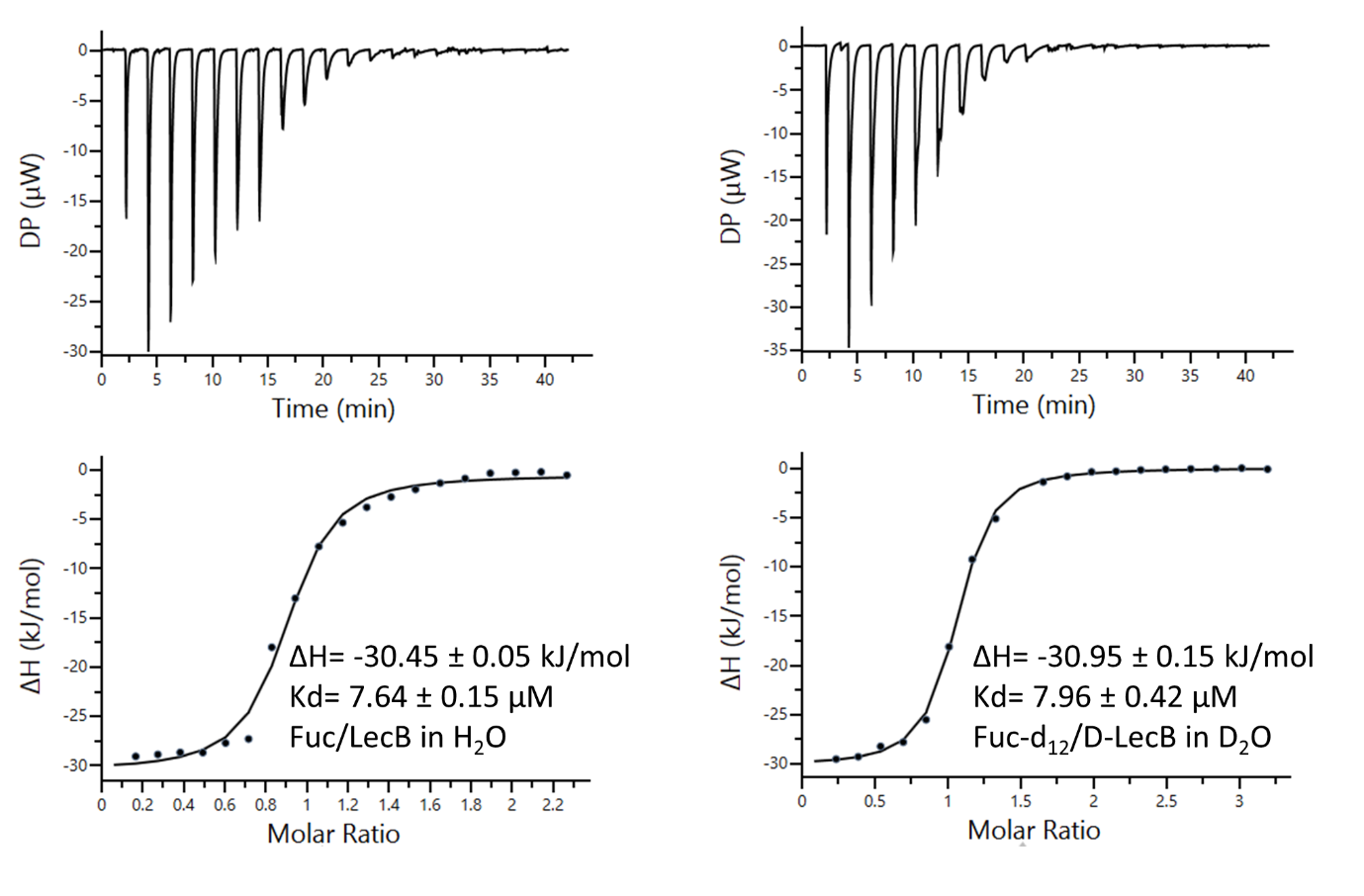

**Figure S4: Isothermal titration calorimetry results of LecB/L-Fuc and D-LecB/L-Fuc-d_12_ binding.** Top: data obtained from 20 automatic injections, 2 μL each, of L-fuc (4 mM)/L-Fuc-d_12_ (7.6 mM) into the cell containing LecB (0.46 mM)/D-LecB (0.50 mM). Lower: plot of the total heat released as a function of total ligand concentration for the titration shown above. The solid line represents the best least-squares fit for the obtained data.

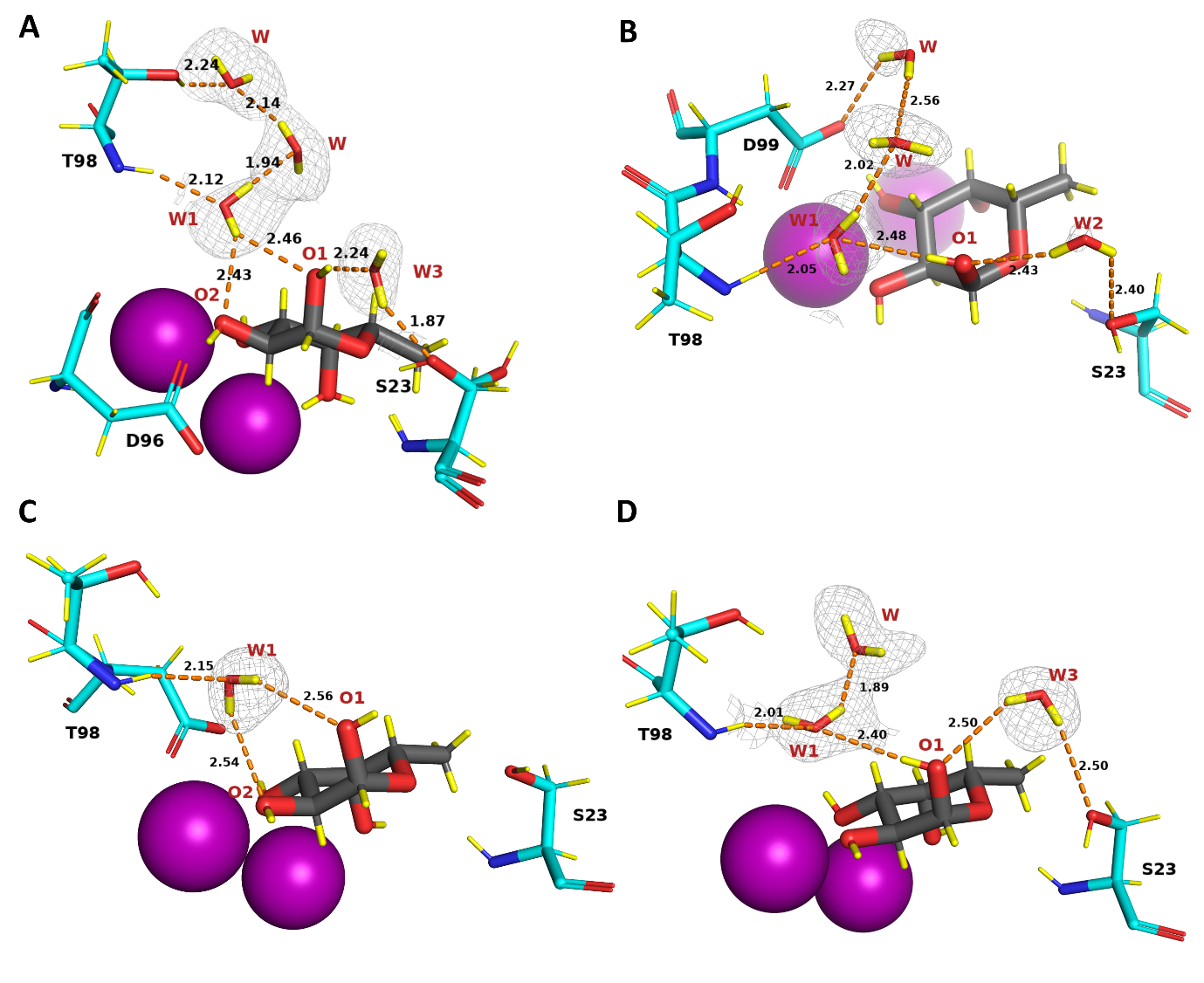

**Figure S5:** **Stick representation of the fucose-binding sites in chains A, B, C and D of the perdeuterated LecB/fucose tetrameric complex.** The 2m*F*_o_-D*F*_c_ neutron density map (grey mesh) is contoured at 0.7σ and is showing highly ordered and contacting water molecules in the binding sites. Hydrogen bonds are shown as orange dashed lines with distances in Å. Fucose is depicted as grey sticks. Deuterium atoms are coloured yellow. Calcium ions are shown as purple spheres.
